# Acute irradiation alters the heterogeneity among medullary thymic epithelial cells

**DOI:** 10.1101/2023.04.03.535248

**Authors:** Kenta Horie, Kano Namiki, Maki Miyauchi, Tatsuya Ishikawa, Mio Hayama, Yuya Maruyama, Takahisa Miyao, Shigeo Murata, Nobuko Akiyama, Taishin Akiyama

## Abstract

The thymus has the ability to regenerate from acute injury caused by radiation, infection, and stressors. In addition to thymocytes, thymic epithelial cells in the medulla (mTECs), which are crucial for T cell self-tolerance by ectopically expressing and presenting thousands of tissue-specific antigens (TSAs), are damaged by these insults and recover thereafter. However, given recent discoveries on the high heterogeneity of mTECs, it remains to be determined whether the frequency and properties of mTEC subsets are restored during thymic recovery from radiation damage. Here we demonstrate that acute total body irradiation with a sublethal dose induces aftereffects on heterogeneity and gene expression of mTECs. Single-cell RNA-sequencing (scRNA-seq) analysis showed that irradiation reduces the frequency of mTECs expressing AIRE, which is a critical regulator of TSA expression, 15 days after irradiation. In contrast, transit-amplifying mTECs (TA-mTECs), which are progenitors of AIRE-expressing mTECs, and Ccl21a-expressing mTECs, were less affected. Interestingly, detailed analysis of scRNA-seq data suggested that the proportion of a unique mTEC cluster expressing Ccl25 and high level of TSAs was severely decreased by irradiation. Overall, we propose that acute irradiation disturbs the heterogeneity and properties of mTECs, and may thereby impair TSA expression for thymic T cell selection.

## Introduction

The thymus provides an environment for development of T cells. Thymic epithelial cells (TECs) are essential for development and selection of thymic T cells by presenting self-antigens and expressing some cytokines and signaling molecules (Abramson and Anderson, 2017). TECs are largely divided into two types based on localization: cortical TECs (cTECs) and medullary TECs (mTECs). cTECs are critical for proliferation and differentiation of early progenitors into CD4^+^CD8^+^ (DP) thymocytes and they also positively select DP thymocytes recognizing a complex of self-peptide and major histocompatibility complex (MHC) with moderate affinity, thereby leading to differentiation of DP thymocytes into CD4^+^CD8^-^ (CD4SP) or CD4^-^CD8^+^ (CD8SP) thymocytes. Subsequently, CD4SP and CD8SP thymocytes interact with mTECs, which ectopically express and present self-peptides derived from tissue-specific antigens (TSAs). CD4SP and CD8SP thymocytes interacting with TSA peptides with high affinity undergo negative selection, or in the case of CD4SP thymocytes, differentiate into regulatory T cells. Autoimmune regulator (AIRE) is a transcriptional regulator that controls TSA expression in mTECs (Anderson et al., 2002). Because dysfunctional mutations of the *AIRE* gene cause onset of human autoimmune disease (Bruserud et al., 2016), it is most likely that TSA expression in mTECs is essential to suppress onset of autoimmune diseases (Derbinski et al., 2001).

Recent studies using single-cell RNA sequencing analysis (scRNA-seq) revealed a high degree of TEC heterogeneity in humans and mice (Bornstein et al., 2018; Dhalla et al., 2020; Park et al., 2020; Wells et al., 2020; Miyao et al., 2022). Bornstein et al. classified mTECs into four subsets (Bornstein et al., 2018). mTEC IV (thymic tuft cells) showing gene expression and chromatin structure similar to intestinal tuft cells have been reported (Bornstein et al., 2018; Miller et al., 2018), in addition to mTEC I, mTEC II, and mTEC III subsets that corresponding primarily to Ccl21a^+^mTEC^lo^, Aire^+^mTEC, and post-Aire mTEC subsets, respectively. Michelson et al. investigated heterogeneity of post-Aire mTECs in more detail. Post-Aire mTECs are an assembly of various mimetic cells harboring signature genes expressed by extra-thymic cell types (Michelson et al., 2022). Several studies using scRNA-seq have suggested a unique mTEC cluster expressing a high level of cell cycle-related genes (Dhalla et al., 2020; Wells et al., 2020; Miyao et al., 2022). A further study demonstrated that this cluster is equivalent to transit-amplifying cells (Miyao et al., 2022), which are proliferative progenitors connecting stem cells and mature cells (Hsu et al., 2014).

Cytoreductive therapies such as chemotherapy and irradiation are frequently used in cancer treatment. Irradiation causes acute involution of the thymus (Kinsella and Dudakov, 2020; Duah et al., 2021). Thymic involution by irradiation is ascribed to severe and acute reduction of thymocytes, mainly of DPs. In addition to thymocytes, TECs are affected by irradiation. Notably, after this acute thymic involution, both thymocytes and TECs are homeostatically recovered (Dudakov et al., 2012; Kaneko et al., 2019; Kinsella and Dudakov, 2020; Duah et al., 2021). Some studies have provided mechanistic insights on the recovery of TECs and the resulting recovery of the thymic organ (Dudakov et al., 2012; Wertheimer et al., 2018; Kinsella and Dudakov, 2020; Duah et al., 2021). Several cytokines and growth factors were reportedly involved in the thymic regeneration. Dudakov et al. have shown that IL-22 from type 3 ILCs is required for TEC proliferation after irradiation (Dudakov et al., 2012). Wertheimer et al. showed that bone morphogentic protein 4 (BMP4) is expressed in thymic endothelial cells and promotes thymic recovery after irradiation (Wertheimer et al., 2018). Mechanistically, BMP4 signaling in TECs causes increased expression of Foxn1, which is critical for TEC development and maintenance. In an earlier study, it was reported that keratinocyte growth factor (KGF) contributes to the thymic recovery from irradiation by enhancing the proliferation and function of thymic epithelial cells (Rossi et al., 2007a). Mechanistically, KGF signaling up-regulates the expression of several target genes necessary for TEC regeneration, such as BMP4. KGF activates the p53 pathway for transcriptional activation. Indeed, the transcription factor p53, which is activated by radiation-induced DNA damage, promotes the regeneration of medullary TECs (mTECs) after irradiation (Rodrigues et al., 2017). In contrast to these positive regulators, TGF-β signaling in TECs inhibits thymic recovery in the early phase after irradiation (Hauri-Hohl et al., 2008).

Although the reduction of TEC frequency by irradiation was known, its effect on TEC heterogeneity and their individual phenotype was not fully characterized. In this study, we performed scRNA-seq analysis of TECs in murine thymus 15 days after sublethal total body irradiation. Data analysis showed that irradiation affects proportions of mTEC subsets. Pseudotime analysis of scRNA-seq data and BrdU decay kinetic analysis have suggested that the decay of Aire^+^mTECs was not affected, but its differentiation might be retarded by irradiation. Moreover, scRNA-seq suggested a selective decrease of an mTEC subset expressing *Ccl25* and a high level of TSAs in addition to a reduction in TSA expression level. Consequently, these data suggested that irradiation affects phenotypes and heterogeneity of mTECs, which may influence thymic selection.

## Results

### mTEC composition was disturbed after irradiation

Because influences of radiation may be altered in different breeding environments, ages, genders, and genetic backgrounds, in addition to the radiation dose, we first detailed recovery dynamics of total thymic cells and TECs (EpCAM^+^ CD45^-^ TER119^-^ cells) from female mice on a 7-week-old C57BL/6 background at several time points after irradiation with a sublethal dose of γ-rays (5.5 Gy) (Figure 1A). Based on binding of UEA-1 lectin and expression level of Ly51, TECs were separated into subpopulation of mTECs (UEA-1^+^Ly51^-^ TECs) and cTECs (UEA-1^-^ Ly51^+^ TECs). As in previous studies (Kaneko et al., 2019), the cell number of thymocytes decreased until Day 4, and then increased to almost the same number on Day 10 as on Day 0 (Figure 1B). In contrast, the number of TECs decreased until Day 6 (Figure 1C). Moreover, the maximum recovery of TECs occurred on Day 15, but still remained below the level on Day 0 (Figure 1C). These recovery dynamics pertain to mTECs, but not to cTECs (Figure 1D and E). Thus, these data indicate that radiation mainly causes a reduction of mTEC cellularity under these conditions.

**Figure 1.**
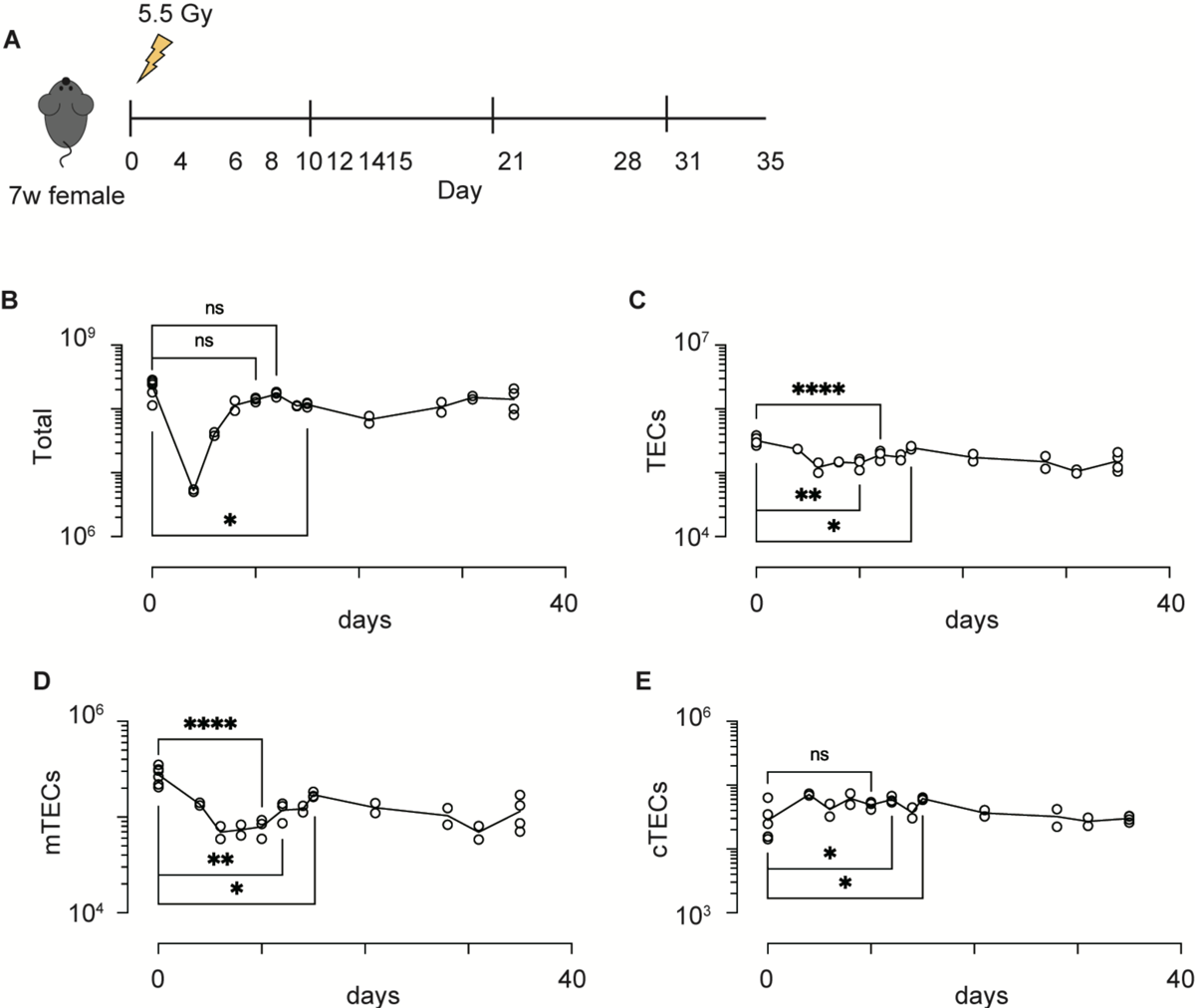
Acute sublethal irradiation caused a reduction and subsequent recovery in cellularity of thymic cells and thymic epithelial cells. **(A)** A schematic diagram of chase experiments of thymic cellularity after irradiation. Seven-week-old female mice with a C57BL/6 background were exposed to sublethal dose radiation. After the indicated days, total thymic cells were analyzed by flow cytometric analysis (**(B-F)** Dynamics in cellularity were evaluated for total thymic cells **(B)**, TECs **(C)**, mTECs **(D)**, and cTECs **(E)** by flow cytometric analysis. Typical profiles of flow cytometric analysis were exhibited in Supplementary Figure 1. *: p < 0.05; **: p < 0.01; ****: p < 0.0001; ns: p > 0.05; Student’s t-test.

Interaction between TECs and thymocytes is critical for development and regeneration of the thymus (Duah et al., 2021). Because mTEC development is dependent on TNF cytokine family signaling from CD4SP thymocytes (Hikosaka et al., 2008; Irla et al., 2008), we detailed aftereffects of radiation on CD4SP thymocytes. CD4SP and CD8SP thymocytes among thymic naïve T cells were separated according to the gating strategy described in a previous study (Xing et al., 2016) (Figure 2A and Supplementary Figure 3). After irradiation, CD4SP thymocyte numbers recovered by Day 15, whereas numbers of DP and CD8SP thymocytes were slightly reduced. CD4SP thymocytes are divided into three sub-populations at different maturation stages, based on expression levels of H2-kb and CD69 (Xing et al., 2016) (Figure 2B). Thus, CD69^hi^ H2-kb^lo^ cells and semi mature cells (SM) are the most immature types of CD4SP thymocytes. CD69^lo^ H2kb^hi^ cells, type 2 mature cells (M2) are the most mature cells, and CD69^hi^ H2kb^hi^ cells, type 1 mature cells (M1), are considered an intermediate differentiation stage between SM and M2 cells (Xing et al., 2016). Ratios of SM in CD4SP were decreased and those of M1 and M2 were increased in the thymus on Day 15. Cell number was slightly decreased for SM1, but not changed for M1 and M2. Thus, development of CD4SPs had largely recovered by Day 15, implying that the reduction of mTECs may not simply be due to reduced cellularity of CD4SP thymocytes.

**Figure 2.**
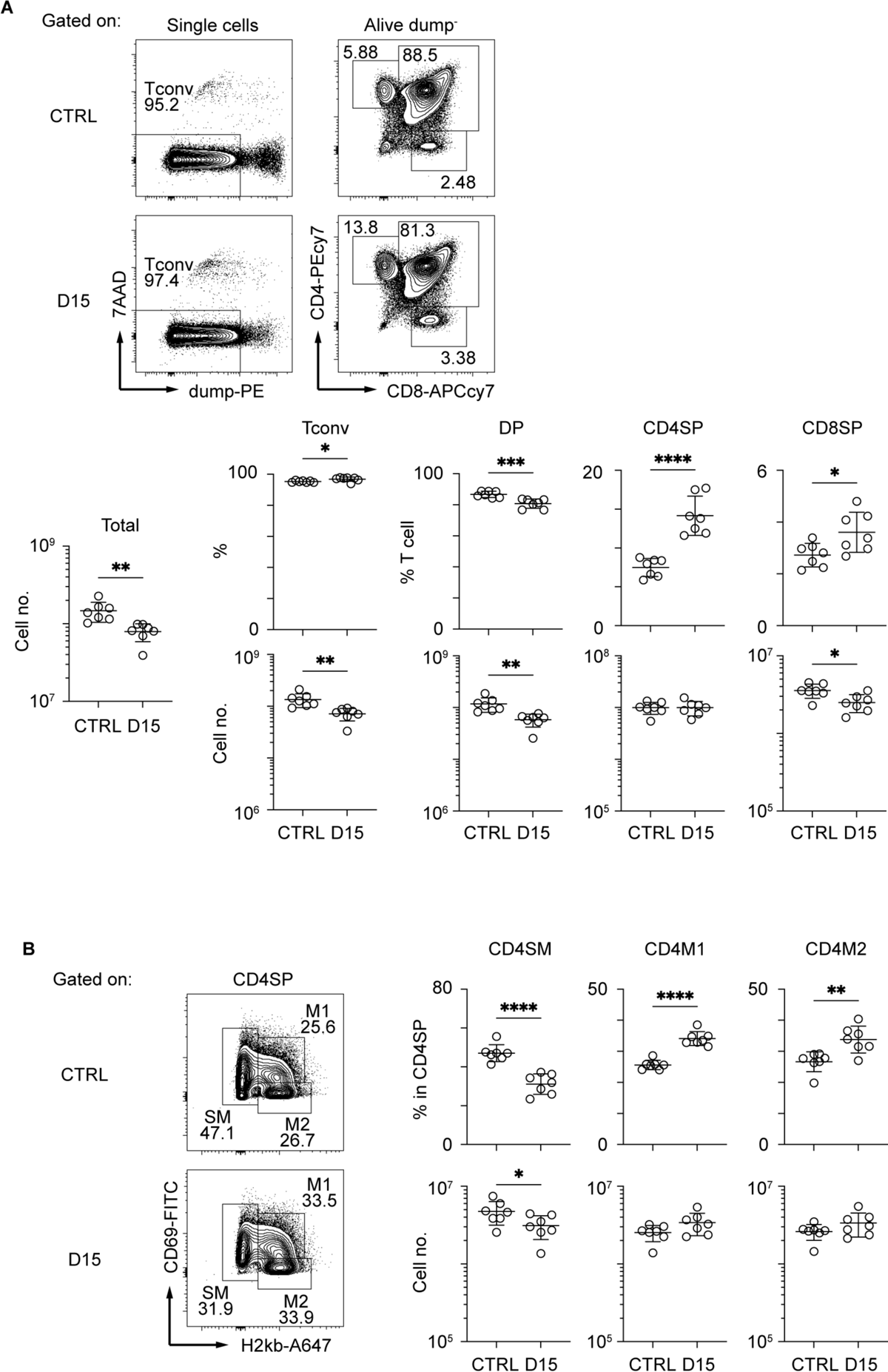
Thymocytes in the medulla were mostly recovered 15 days after sublethal irradiation. **(A)** Representative flow cytometric profiles of thymocytes. Dump^-^ cells are defined as 7AAD^-^ CD25^-^ CD44^-^ γδTCR^-^CD1d-tetramer^-^ cells. Data are summarized in figures below. Tconv indicates Dump-cells. DP, CD4SP, and CD8SP indicates CD4^+^CD8^+^Dump^-^, CD4^+^CD8^-^Dump^-^, and CD4^-^CD8^+^Dump^-^ cells ****: **: p < 0.05; **: p < 0.01; ***: p < 0.001; ****: p < 0.0001; two-tailed t test.

### Differentiation stages from TA-mTECs to Aire^+^mTEC were affected 15 days after irradiation

We next sought to find effects of irradiation on heterogeneity and transcriptomics of TECs. We performed scRNA-seq of TECs from mice 15 days after irradiation and age-matched control mice because the number of TECs and mTECs was largely recovered by this time. TECs (CD45^-^TER119^-^EpCAM^+^ cells) were sorted from collagenase-digested thymic cells of these mice for scRNA-seq. To normalize individual variation, TECs from 3 mice were pooled for each condition. TEC data from irradiated and control mice were integrated and compared using the Seurat package (Stuart et al., 2019). After filtering out cells with poor-quality expression data, 8,303 cells were used for downstream analysis. Among selected cells, 5,070 cells were irradiated TECs and 3233 cells were control TECs. Unsupervised graph-based clustering and UMAP dimension reduction of scRNA-seq data separated TECs into 16 clusters (Figure 3A). Based on expression levels of TEC marker genes (Figure 3B, C and Supplementary Figure S2A), we assigned clusters 0, 4, 5 and 11 to *Ccl21a*^+^ mTECs, clusters 1 and 7 to *Aire*^+^ mTECs, cluster 10 to tuft-like mTECs, clusters 12 and 14 to Post-Aire mTECs, cluster 8 to cTECs, cluster 13 to nurse cTECs, and cluster 3 to probable Pdpn^+^ TECs (Onder et al., 2015). TA-mTECs are concordant with clusters 6 and 9, which are separated most likely because of differences in cell cycle phase proportions (Supplementary Figure S2C, D and E). Cluster 2 seemed to contain both *Aire*^+^ and *Aire*^-^ TECs (Figure 3B and C) and its average expression level of *Aire* was lower than that of cluster 1; therefore, it was assigned to Late-Aire mTECs (Ferreirinha et al., 2021).

**Figure 3.**
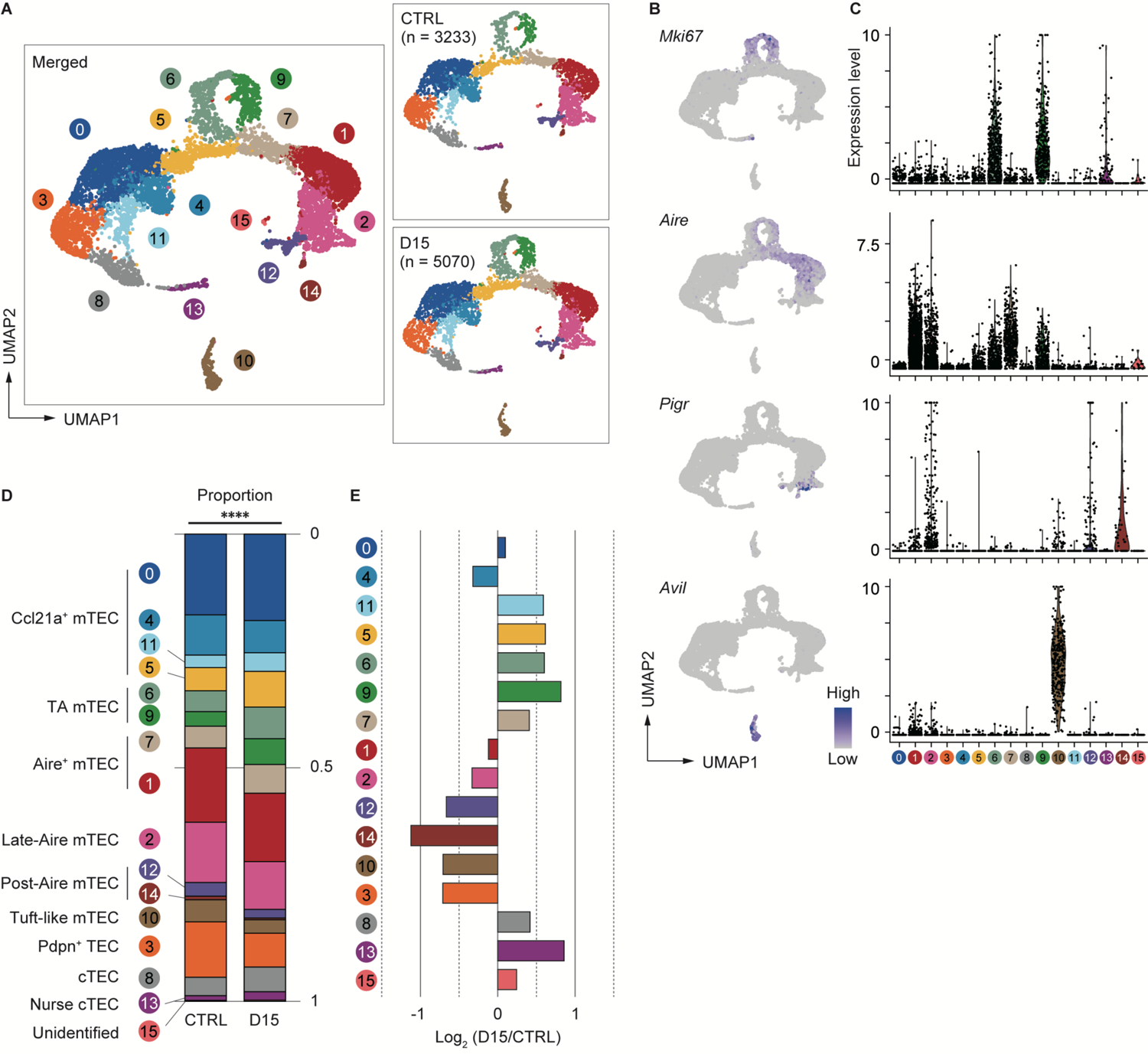
Heterogeneity of mTECs remained altered 15 days after sublethal irradiation. **(A)** UMAP plots of scRNA-seq data of TECs from mice 15 days after irradiation (D15) and age-matched control mice (CTRL). After being integrated to eliminate a batch effect, two datasets are split (right figures). Parentheses indicate cell numbers used for analysis after filtering to remove low-quality data. Cluster numbers are indicated. **(B)** Heatmaps of marker gene expression (counts per 10^4^ counts) projected onto UMAPs. **(C)** Violin plots of marker gene expression (scaled counts per 10^4^ counts) **(D)** Stacking bar plots of proportions of each cluster in TECs from mice 15 day after irradiation (D15) and age-matched control mice (CTRL). ****: p < 0.0001; Chi-square goodness of fit test. **(E)** Bar plots of log2-transformed changes in proportions of TEC clusters in mice 15 day after irradiation (D15) and age-matched control mice (CTRL). Wilcoxon rank sum test. SST: steady-state thymus; PIT: post-irradiation thymus

Proportions of TEC subsets were compared between the two conditions and log2-transformed changes of proportions were determined (Figure 3D and E). Chi-square goodness of fit tests showed a significant difference in the proportion of TEC subsets between irradiated and non-irradiated cells(Figure 3D). Proportions of Late-Aire mTEC, Post-Aire mTECs, tuft-like mTECs, and Pdpn^+^TECs were reduced (Figure 3E). In contrast, proportions of CCL21a^+^ mTECs, TA-mTECs, and cTECs were increased. In Aire^+^ mTECs, cluster 1 was slightly decreased. In contrast, cluster 7, which is a transition stage from TA-mTECs to Aire^+^mTECs (Miyao et al., 2022), was increased. Overall, these data suggested that sublethal-dose radiation caused abundant aftereffects in Aire^+^ mTECs and their progeny, i.e. Late-Aire, Post-Aire, and tuft-like mTECs, more than TA-TEC progenitors and CCL21a^+^ mTECs.

We considered two possible explanations for the effect of radiation on the ratio of Aire^+^mTECs and their later differentiation stages. The first is that irradiation may promote a decrease in fully differentiated Aire^+^mTECs, which are post-mitotic and are susceptible to cell death (Gray et al., 2007). The second is that differentiation from TA-mTEC progenitors into Aire^+^mTECs was repressed by irradiation.

A previous report suggested that pseudotime analysis may predict the differentiation rate of cells (Zhou et al., 2022). We then performed single-cell trajectory analysis of TEC scRNA-seq data using the Monocle 3 package (Cao et al., 2019) to determine the pseudotime of each cell. Cells closest to TA-mTECs were chosen as root cells, which is the starting point for the pseudotime trajectory (Figure 4A). To evaluate differences in the differentiation rate, the pseudotime mean of cells around the branching of TA-mTEC, Aire^+^mTEC and Late Aire mTEC stages was calculated for control and irradiated TECs. The pseudotime mean of TECs receiving radiation was significantly lower than that of control TECs (Figure 4B), implying that the differentiation rate of mature mTECs is slowed 15 days after irradiation.

**Figure 4.**
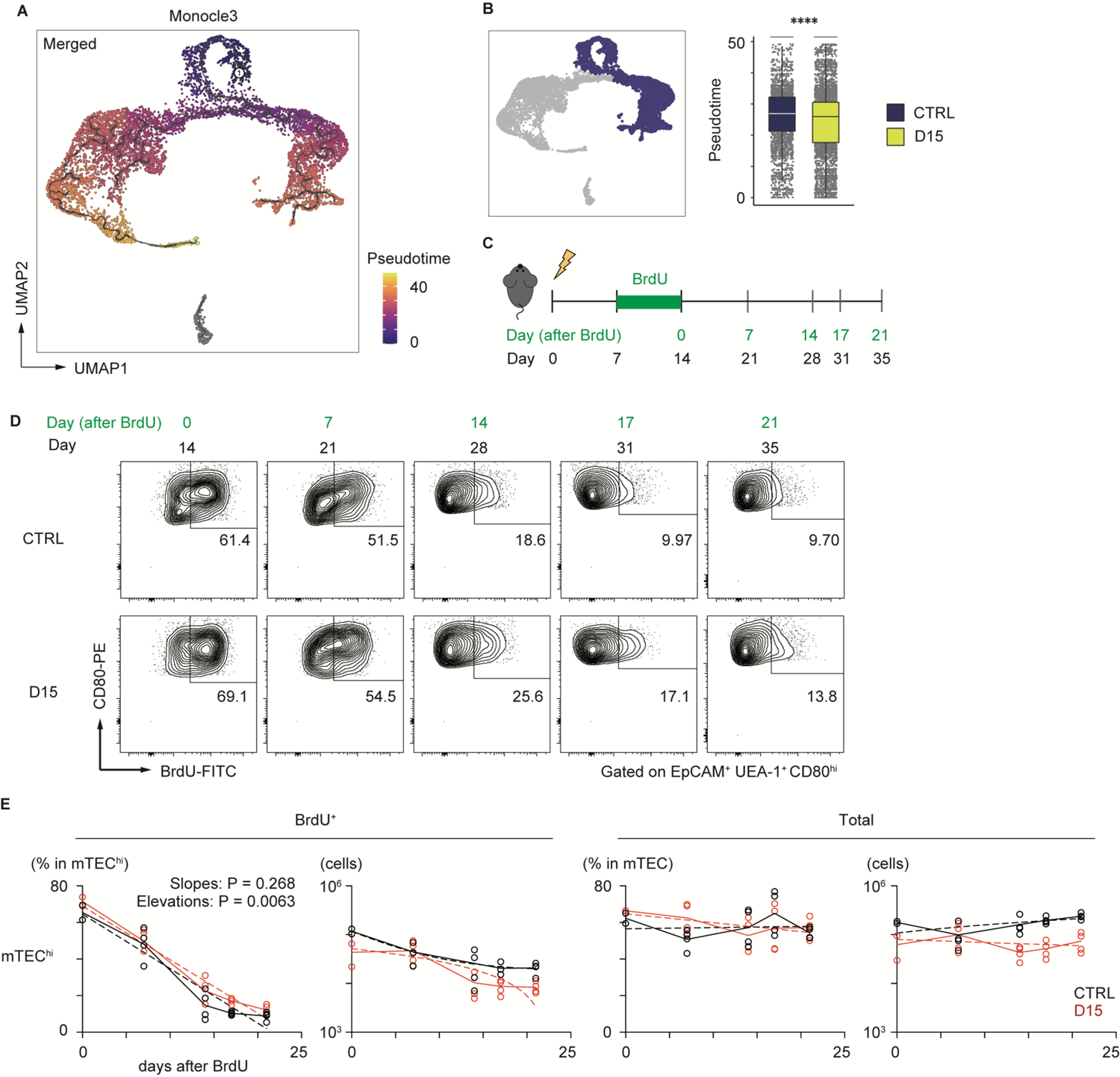
Irradiation did not influence the reduction kinetics of mature mTEC frequency. **(A)** The trajectory graph and pseudotime projected onto the UMAP plot of scRNA-seq data of TECs. **(B)** Box plots show pseudotime means of TA-mTECs, Aire^+^mTECs, and Late-Aire mTECs from mice 15 day after irradiation (D15) and age-matched controls (CTRL). Each dot indicates the pseudotime of an individual cell. ****: p < 0.0001; Wilcoxon rank sum test. **(C)** A schematic diagram of *in vivo* BrdU-labeling experiment. BrdU was supplied in drinking water from day 7 to day 14 after irradiation. **(D)** Representative flow cytometer plots of mTEC^hi^ incorporating BrdU. mTEC^hi^ were defined as CD80^hi^EpCAM^+^ UEA1^+^ CD45^-^ TER119^-^cells **(E)** Decay kinetics of BrdU-labeled cells after the end of BrdU labeling. Solid lines connect means at each timepoint. Broken lines in plots of percentage against days indicate simple linear regression of average values. Broken lines in plots of cellularity were fitted by the least square method.

In order to evaluate reduction kinetics of differentiated mTECs, we performed a 5-bromo-2’-deoxyuridine (BrdU)-labeling assay. BrdU labeling of TA-mTECs and differentiated mTECs was performed by supplying BrdU in drinking water during the recovery period (Days 7 to 14 in Figure 4C). After BrdU labeling, the frequency of BrdU-labeled mTECs (UEA-1^+^EpCAM^+^) was monitored by flow cytometric analysis for 3 weeks (Figure 4D). Immediately after the end of the BrdU up-take, the ratio of BrdU-labeled cells in mTECs expressing high levels of CD80 (mTEC^hi^), which include TA-mTECs, Aire^+^mTECs and Post-Aire mTECs, was 69.1% of irradiated mTECs^hi^, versus 61.4% of the control group, suggesting similar proliferation of mTECs^hi^ in the two conditions. As expected, the proportion of BrdU-labeled cells in mTECs^hi^ gradually decreased and plateaued by Day 17 after BrdU-labeling, whereas total TEC numbers were practically constant during this period (Figure 4E). Notably, reduction kinetics of BrdU-labeled mTECs^hi^ were similar between the two conditions (Figure 4E). Thus, while it is possible that mTECs^hi^ is more susceptible to apoptosis immediately after irradiation, these results suggest that the sensitivity of mTECs^hi^ to cell death might not be largely different between irradiated and non-irradiated thymus at 15 days post-irradiation. Accordingly, it appears that the reduction in the frequency of mTECs^hi^ at Day 15 is likely due to a repression in mTEC differentiation during the recovery period, and/or preferential cell death of mTECs^hi^ early after irradiation.

### Expression levels of tissue-specific antigens were altered by irradiation

We next investigated irradiation-induced aftereffects in gene expression of mTECs, using scRNA-seq data. We focused on gene expression changes in Aire^+^mTECs and their later differentiation stages because of their greater susceptibility to radiation. *Aire* expression was highest in Aire^+^ mTECs, comprising clusters 1 and 7, and moderate in Late-Aire mTECs (cluster 2) and TA-mTECs comprising clusters 6 and 9 in control mice (Figure 5A). Radiation caused a slight down-regulation of *Aire* expression in TA-mTECs and up-regulation of Late-Aire mTECs at Day 15, whereas *Aire* expression was not significantly altered in Aire^+^mTECs (Figure 5A).

**Figure 5.**
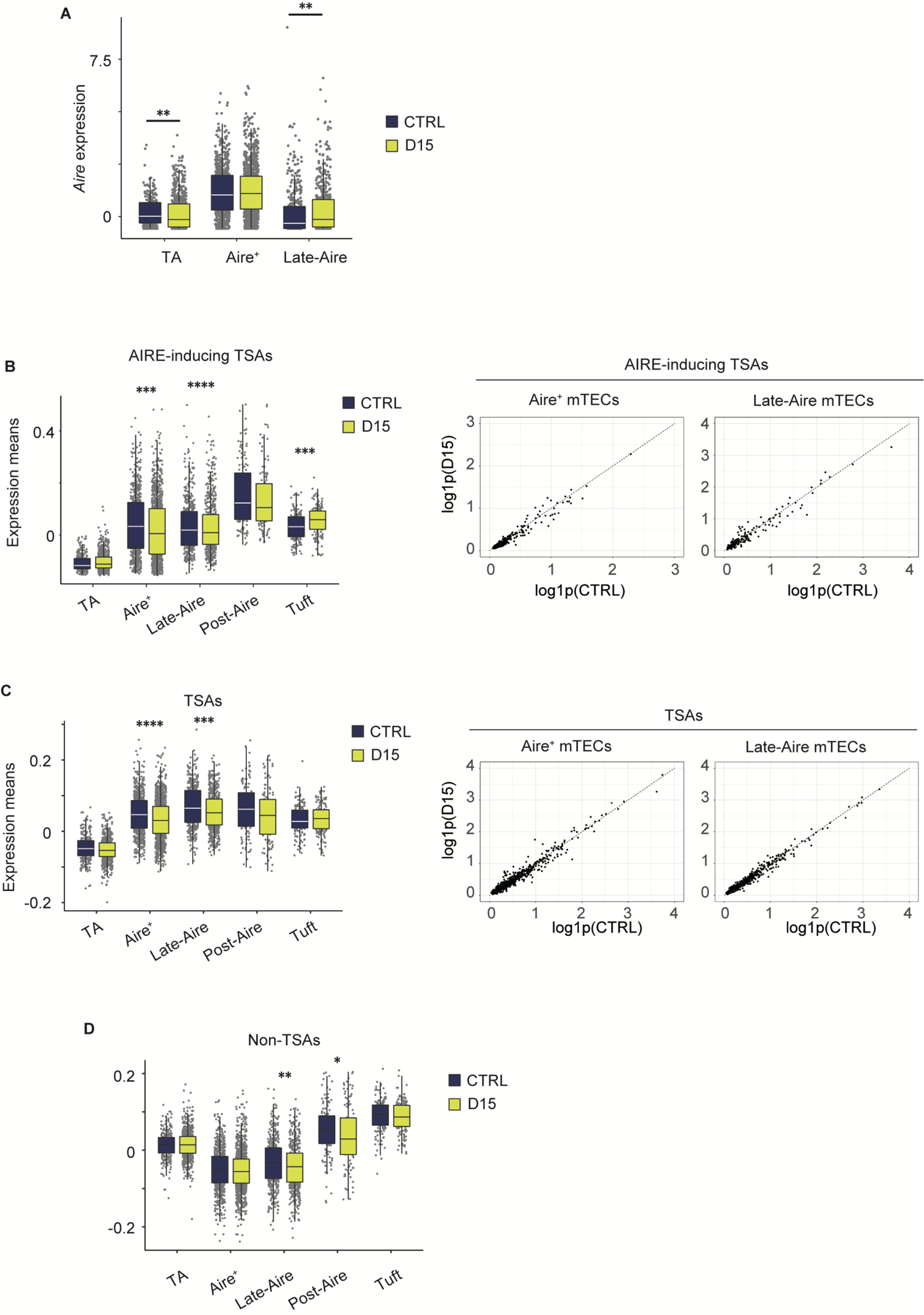
Expression level of AIRE and tissue-specific antigen genes were altered 15 days after acute sublethal irradiation. **(A)** A box plot of means of *Aire* expression in indicated cell subsets (TA, TA-mTECs, Aire^+^, Aire^+^mTECs, and Late-Aire, Late-Aire mTECs) from mice 15 days after sublethal irradiation (D15) and controls (CTRL). **(B-D)** Box plots for mean TSA expression, AIRE-inducing TSAs and non-TSAs from mice 15 days after irradiation (D15) and controls (CTRL). Scatter plots show log-transformed expression values of AIRE-inducing TSAs (B) or TSAs (C) in Aire^+^ and Late-Aire mTECs. Log1p indicates ln(1 + *x*); the natural logarithm of 1 + *x*. TA, Aire^+^, Late-Aire and Tuft indicate TA-mTECs, Aire^+^mTECs, Late-Aire mTECs, and tuft-like mTECs. TSA and Aire-inducing TSAs were selected according to a previous study (Sansom et al., 2014). *: p < 0.05; **: p < 0.01; ***: p < 0.001; ****: p < 0.0001; Wilcoxon rank sum test. For gene expression analysis, genes with more than 500 UMI counts were selected as practically expressed genes.

Because Aire expression was differentially altered, we next investigated aftereffects on TSA expression in each subset. A list of TSAs and AIRE-inducing TSAs (Supplementary Table 1), which are down-regulated by *Aire* gene deletion (Sansom et al., 2014), were used for this analysis. Mean expression levels of all TSAs and AIRE-inducing TSAs were slightly reduced in Aire^+^mTECs and Late-Aire mTECs from irradiated mice, compared to control mice (Figure 5B and C) although changes in individual TSA expression level are small (Figure 5B and C). Interestingly, for Late-Aire mTECs, Non-TSA genes in addition to TSA genes were reduced (Figure 5D), suggesting a possible effect of radiation on general gene expression mechanisms. For tuft-like mTECs, total TSA expression was unchanged and the mean expression level of AIRE-inducing TSAs was increased (Figure 5B and C). Interestingly, although Aire expression was unchanged in Aire^+^ mTECs and increased in Late-Aire mTECs (Figure 5A), mean expression levels of AIRE-inducing TSAs were decreased in both mTECs. Thus, the change in expression level of AIRE-inducing TSAs by irradiation was not correlated with the change in AIRE expression level, implying that other regulatory mechanism(s) controlling ectopic TSA expression might be disturbed by irradiation.

### mTECs expressing Ccl25 were selectively decreased 15 days after irradiation

We next conducted a detailed analysis of Aire^+^mTECs and Late-Aire mTECs, which are considered mature mTECs expressing high levels of MHC class II and co-stimulatory molecules. Because the TSA expression profile is altered by irradiation, we performed subclustering and UMAP dimension reduction based on TSA expression levels for these mTECs. We found that Aire^+^mTECs and Late-Aire mTECs are separated into 6 subclusters by expression levels of TSAs (Figure 6A). Embedding these subclusters in the original UMAP plot suggested that subclusters 0 and 2 are mainly concordant with Aire^+^mTECs and that subclusters 1, 3, 4, and 5 are concordant with Late-Aire mTECs (Figure 6B). Comparisons of the proportions of these subclusters between irradiated TECs and controls revealed that subcluster 4 was most severely decreased by irradiation (Figure 6C and D). Subcluster 4 is located close to the border between Aire^+^mTECs and Late-Aire mTECs in the original UMAP plot (Figure 6B). Consistently, Monocle 3 analysis suggested that the average pseudotime of cluster 4 cells is between that of cluster 0 and cluster 1 (Figure 6E and Supplementary Figure S2F), which are typical clusters of Aire^+^mTECs and Late Aire mTECs, respectively. Moreover, expression of Aire in subcluster 4 is intermediate between those of clusters 0 and 1 (Figure 6F). Thus, this radiation-sensitive subcluster 4 most likely corresponds to an intermediate subset between Aire^+^mTECs and Late-Aire mTECs.

**Figure 6.**
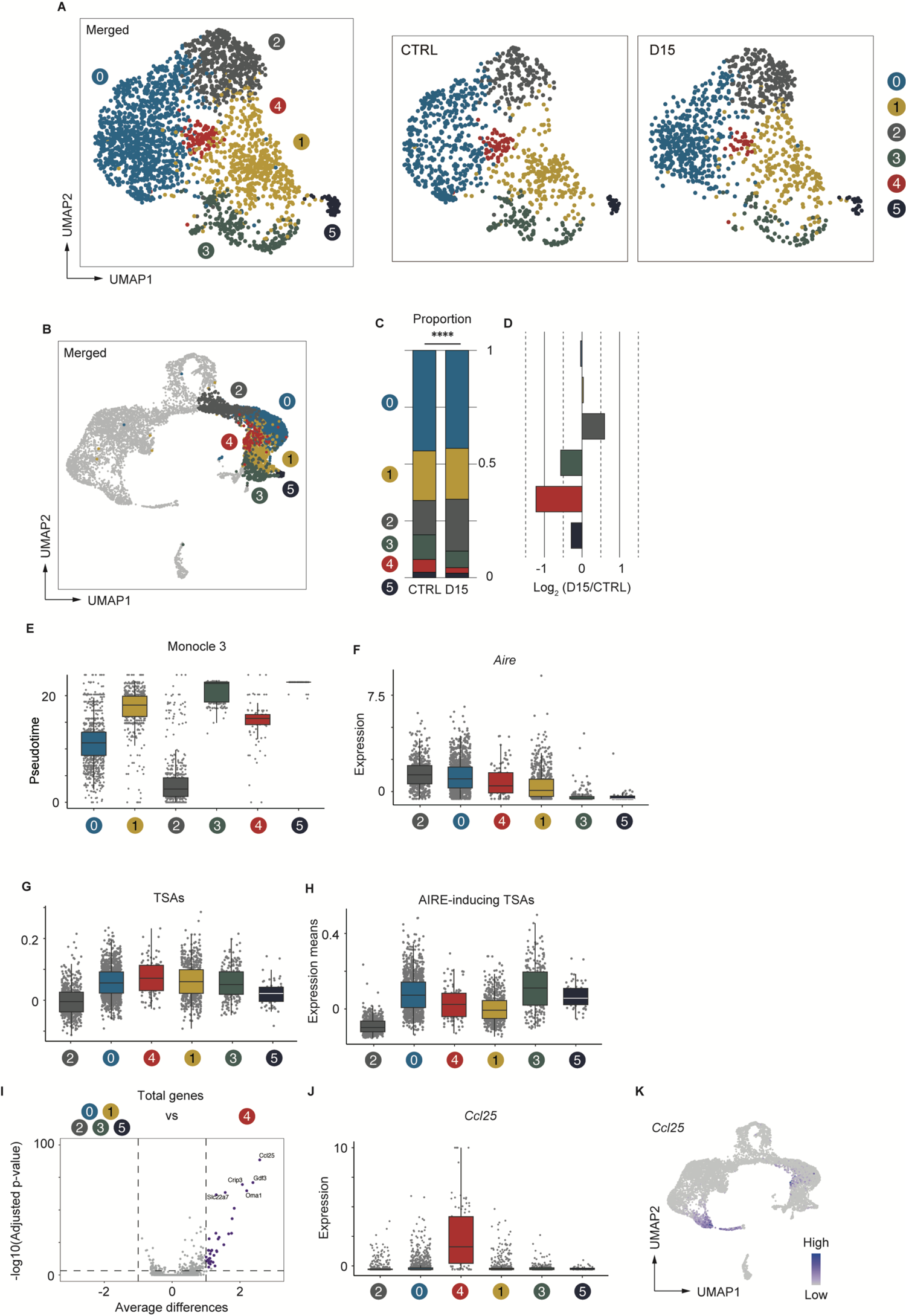
*Aire*^+^ mTECs expressing CCL25 and a high level of TSAs were preferentially decreased 15 days after acute irradiation. **(A)** UMAP plot for subclustering analysis of Aire^+^mTECs and Late-Aire mTECs. Only TSA genes were used for subclustering and UMAP reduction. UMAP plot of D15 is shown after downsampling to equalize the number of cells in the CTRL. **(B)** Embedding clusters of (A) in the original UMAP plot using all cells and genes. **(C)** Stacking bar plots of proportions of each subcluster from TECs mice 15 day after sublethal irradiation (D15) and controls (CTRL). **(D)** Bar plots of log2-transformed changes in proportions of TEC subclusters from mice 15 day after irradiation (D15) and age-matched control mice (CTRL) **(E)** Box plots shows pseudotimes of cells in subclusters determined based on TSA expression. Each dot indicates a pseudotime of an individual cell. Box plots of *Aire* expression **(F)**, means of AIRE-inducing TSA expression **(G)**, and means of TSA expression **(H)** in each subcluster are indicated. **(I)** A volcano plot of changes in gene expression between subcluster 4 and other subclusters. Purple dots indicate significantly upregulated genes in subcluster 4 (p-value < 0.05, average differences > 1). P-values were calculated with the Wilcoxon rank-sum test with Bonferroni correction. **(J)** Box plots of *Ccl25* expression in each subcluster. **(K)** A heatmap of Ccl25 expression projected onto UMAP. Gene expression values were scaled counts per 10^4^ counts.

Because subcluster 4 was detected by subclustering and UMAP analysis based on TSA expression, we analyzed TSA expression in subcluster 4. Interestingly, expression levels of TSAs are highest among these subclusters. However, expression of Aire-inducing TSAs was moderate and lower than that of subcluster 0. This finding implies that the profile of TSAs expressed in mature mTECs is dynamically changed during maturation. We next sought to find marker genes specific to subcluster 4. Differential gene expression analysis of subcluster 4 and other subclusters revealed significant up-regulation of several genes in cluster 4 (Figure 6I). In particular, C-C motif chemokine ligand 25 gene (*Ccl25*) is most differentially expressed in cluster 4 (Figure 6I and J). Thus, the analysis suggested that, in addition to cTECs, a specific subset of Aire^+^mTECs highly expresses CCL25 (Figure 6K). Importantly, these CCL25-expressing mTECs may be decreased after acute irradiation.

## Discussion

In this study, we investigated aftereffects of sub-lethal dose irradiation on murine TEC phenotypes. Data suggested that even if the number of total thymic cells recovered, recovery of mTEC cellularity was not complete. scRNA-seq analysis suggested that reduction occurs in Aire^+^mTECs and mTECs in later differentiation stages, which are considered mature mTECs. Importantly, we found that the frequency of TA-mTEC progenitors was less influenced by radiation, compared to mature mTECs. Label retention experiments suggested that the reduction in cellularity of mature mTEC subsets was not due to enhanced mTEC cell death, but may be due to decelerated differentiation from TA-mTECs to Aire^+^mTECs. The important question is what mechanisms of differentiation are affected by irradiation.

TNF family RANKL signaling is critical for differentiation of Aire^+^mTECs (Rossi et al., 2007b; Akiyama et al., 2008; Hikosaka et al., 2008). Moreover, a previous study revealed that RANKL is up-regulated in CD4SP and type 3 ILCs after bone marrow transplantation with irradiation, and the administration of recombinant RANKL protein boosts the regeneration of mTECs (Lopes et al., 2017). On the other hand, a recent study suggested that RANKL signaling may promote differentiation from immature mTECs to TA-mTECs (Wells et al., 2020). It should be noted that the ratio of TA-mTECs was higher than that of controls. Accordingly, it is unlikely that impairment of RANKL signaling is solely responsible for reduction of Aire^+^ mTECs and their descendants. This idea may be supported the fact that CD4SP cells were largely rescued 15 days after irradiation. Thus, it is an open question to identify the mechanism by which irradiation causes a selective effect on development of these mTECs.

One previous study revealed a critical role of IL-22 for TEC regeneration after irradiation (Dudakov et al., 2012). IL-22 is expressed in type 3 innate lymphoid cells via signaling of IL-23, expressed in dendritic cells sensing the reduction of CD4 CD8 double-positive thymocytes. This mechanism promotes proliferation of TECs in relatively early stages of thymic recovery. Thus, it is likely that the IL-22-dependent event occurs ≤15 days after irradiation. However, given that IL-22 signaling enhances proliferation of TECs, most likely including proliferative TA-mTECs, it may be that TA-mTECs receiving IL-22 signaling are less able to differentiate into Aire^+^ mTECs. Reduction of Aire expression in TA-mTECs after irradiation suggests that properties of TA-mTECs can be changed by irradiation. To test this hypothesis, it is necessary to compare phenotypes of irradiated and non-irradiated TA-mTECs in detail. However, presently, it is difficult to discriminate between TA-mTECs and Aire^+^mTECs by flow cytometry. Identification of a cell surface marker for TA-mTECs will help overcome this problem.

One surprising finding in our scRNA-seq analysis is that an mTEC subset expressing high levels of CCL25 is present in the Late-Aire mTEC cluster. CCL25 is highly expressed in cTECs and controls recruitment of T cell progenitors derived from bone marrow into the thymus (Wurbel et al., 2000). Interestingly, a recent study suggested that CCL25 is directly regulated by AIRE (Nishijima et al., 2022). Moreover, the same authors reported that CCL25-deficient mice showed autoimmunity in some organs, which was similar to the phenotype of *Aire*-deficient mice. Therefore, in addition to detailed characterization, an important future study should address whether CCL25-expressing mTECs are critical to suppress onset of autoimmunity.

One limitation of this study is that only one experiment was conducted for scRNA-seq analysis of irradiated and control samples. Therefore, efforts were made to minimize individual differences among mice by pooling TECs from three mice. Furthermore, a comparison of the integrated data under these two conditions suggests that the batch effect due to experimental error is small. However, future validation of the present scRNA-seq data is needed through flow cytometry analysis of cell subsets with appropriate cell surface markers and RNA-seq analysis of sorted cell subsets.

Several studies have suggested that radiation increases the risk of autoimmune diseases, such as diabetes and thyroid autoimmune disease (Schaue, 2017; Yahyapour et al., 2018). Most studies of irradiation-induced autoimmune diseases have focused on lymphocytes. In contrast, our findings imply that influence on mTECs may contribute to increased risk of autoimmune diseases due to irradiation during radiotherapy and disasters. Indeed, a previous study suggested that the recovery of mTEC numbers was delayed compared to mature thymic T cells after bone marrow transplantation in mice, leading to autoimmunity (Alawam et al., 2022). Our data also suggested that the percentage of mature mTECs in the total TECs decreased 15 days after the total-body irradiation, despite the recovery of CD4SP cells. In addition to the decrease in the frequency of mature mTECs, the expression level of TSA in mature mTECs was also decreased. Thus, these two effects on mTECs may additively increase the risk of autoimmunity due to the failure of the mTEC environment to recover after irradiation.

Several studies have suggested that radiation increases the risk of autoimmune diseases, such as diabetes and thyroid autoimmune disease (Schaue, 2017; Yahyapour et al., 2018)). Whereas most studies have focused on the dysregulation of peripheral lymphocytes, the disturbance of the thymic self-tolerance mechanism may also contribute to the increased risk of autoimmune diseases after exposure to radiation through radiotherapy or disasters. Indeed, a recent study showed that the recovery of mTEC numbers was delayed compared to mature thymic T cells after bone marrow transplantation in mice, leading to autoimmunity (Alawam et al., 2022). O Our findings may further support this, as we observed a decrease in the percentage of mature mTECs in the total TECs 15 days after total-body irradiation, despite the recovery of CD4SP cells. In addition, the expression level of TSA in mature mTECs was also decreased. Thus, these two effects on mTECs lead to the impairment of the thymic medullary environment responsible for self-tolerance after irradiation and may combine to increase the risk of autoimmunity. Further analysis of the variation in the TCR repertoire of mature thymic T cells after irradiation may provide some insight into this issue.

Overall, our study suggests an impact of irradiation on properties and heterogeneity of mTECs, which may disrupt thymic self-tolerance mechanisms, thereby increasing the risk of autoimmune diseases. Further verification of this hypothesis is important for discovering mechanisms underlying increased risk of autoimmune disease onset due to irradiation and other thymic insults.

## Materials and Methods

### Mice

All mice were housed under specific pathogen-free conditions according to Guidelines of the Institutional Animal Care and Use Committee of RIKEN, Yokohama Branch (2018-075). C57BL/6 female mice (Clea, Japan) were used for all experiments. Seven-week-old female mice received 5.5 Gy of γ-irradiation. For scRNA-seq analysis, and flow cytometric analysis of thymocytes, mice were used 15 days after irradiation. For BrdU labeling, 0.8 mg/mL BrdU (Nacalai, Kyoto, Japan) was supplied with sterile drinking water for 7 days.

### Antibodies and reagents

Antibodies and reagents employed included APCCy7-anti-mouse CD45 (BioLegend, clone 30 F-11, Cat#103116), APCCy7-anti-mouse TER119 (BioLegend, clone TER-119, Cat#557853), BV711-anti-mouse EpCAM (CD326) (BioLegend, clone G8.8, Cat#118233), PE-anti-mouse CD80 (eBioscience, clone 16-10A1, Cat#12-0801-81), Alexa647-anti-mouse Ly51 (BioLegend, clone 6C3, Cat#108312), Biotinylated UEA-1 (Vector Laboratories, Cat#B-1065), PECy7-streptavidin (eBioscience, Cat#25-4317-82), PECy7-anti-mouse CD4 (BioLegend, clone RM4-5, Cat#100528), APCCy7-anti-mouse CD8 (BioLegend, clone 53-6.7, Cat#109220), PE-anti-mouse CD44 (BioLegend, clone IM7, Cat# 103007), PE-anti-mouse CD25 (BioLegend, clone PC61, Cat#102007), PE-anti-mouse TCRγ/δ (BioLegend, clone GL3, Cat#118107), FITC-anti-mouse CD69 (BioLegend, clone H1.2F3, Cat#104506), AlexaFluor647-anti-mouse H2-kb (BioLegend, clone AF6-88.5.5.3, Cat#116512), FITC-anti-BrdU (BD, BrdU flow kit, Cat#559619), Zombie Aqua fixable viability kit (BioLegend, Cat#423101), and anti-mouse CD16/32 (BioLegend, clone 93, Cat#101302)

### Isolation and analysis of TECs from mice

Mice were sacrificed with CO_2_ and thymi were dissected. Individual thymi were minced with razor blades and thymic fragments were obtained by removing the supernatant and by pipetting minced thymi up and down in 1 mL RPMI1640 (Wako). Thymic fragments were digested in 0.05U/mL Liberase (Roche) and 0.01%w/v DNaseI (Sigma-Aldrich) in RPMI1640 by incubation at 37°C 3 x 12 min. Supernatant from the digestion was added to the same volume of FASC buffer (2% fetal bovine serum; FBS in phosphate-buffered saline; PBS) containing 1 nM EDTA to stop reactions. The cell suspension was incubated with anti-mouse CD16/32 in FACS buffer to block non-specific binding, and stained with primary antibodies (anti-EpCAM, -CD45, -TER119) in FACS buffer. 7-Amino-Actinomycin D (7-AAD) was added to label dead cells. EpCAM^+^ cells were sorted for scRNA-seq using a FACS Aria instrument (BD). For BrdU detection, cells were stained with primary antibodies (anti-EpCAM, -Ly51, -CD45, -TER119, -CD80, biotinylated UEA-1) in FACS buffer and sequentially incubated with secondary reagent (streptavidin) in FACS buffer following a blocking step. Dead cells were labeled with Zombie Aqua fixable viability kit. Flow cytometry data were analyzed with FlowJo10.

### BrdU-labeling experiment

BrdU up taken in DNA was detected using a BrdU flow kit (BD, Cat#559619). In brief, cells were fixed and permeabilized with Cytofix/Cytoperm buffer after cell surface staining. To expose BrdU up taken by DNA, cells were incubated with 300mg/mL DNaseI in DPBS for 1 h at 37°C before anti-BrdU staining.

### Library preparation for droplet-based single-cell RNA-sequencing

For scRNA-seq analysis, cell suspensions of steady-state thymi or post-irradiated thymi from three mice were prepared and pooled. Cellular suspensions were loaded onto a Chromium instrument (10× Genomics) to generate a single-cell emulsion. scRNA-seq libraries were prepared using Chromium Single-Cell 3′ Reagent Kit v3 Chemistry.

### scRNA-seq analysis of TECs

Library paired-end sequencing was performed on an Illumina HiSeq2000. Output FASTQ files were processed using Fastp, demultiplexed, and aligned to the mm10 reference genome with Cell Ranger v5.0.0. Processing with Cell Ranger was performed on the HOKUSAI supercomputer at RIKEN. Downstream analyses were performed with the Seurat package, version 4. Transcript-by-cell metrics were made from output files and contained genes expressed in at least five cells. Cells with a percentage of mitochondrial transcripts < 15% and 7500 > the number of feature RNA > 2000 were used for further analyses. For normalization, feature counts of each cell were divided by total counts of that cell and were multiplied by 10^4^. The 3000 most variable genes were selected with variance stabilizing transformation to perform principal component analysis (PCA). After integration of two data sets with 2000 anchor features, scaling data, PCA and UMAP, followed by shared-nearest-neighbor graph construction were performed. Shared-nearest-neighbor graph construction was performed with the top 20 PCs and resolution of cell clustering analysis was set at 0.5. Subclustering analysis was performed with only TSA genes defined as a list (Sup table 1). Cluster-based differential gene expression analysis was performed using FindMarkers in the Seurat package with scaled expression values. Pseudotime analysis was performed with the Monocle3 package.

### Isolation and analysis of thymocytes

Mice were sacrificed with CO_2_ and thymi were dissected. Individual thymi were ground with glass slides in 3 mL 5% FBS in RPMI1640 (Wako). Cells were centrifuged, incubated with anti-mouse CD16/32 in FACS buffer to block non-specific binding and stained with primary antibodies (anti-CD4, -CD8, -CD69 and -H2-kb) in FACS buffer. 7-Amino-Actinomycin D (7-AAD) was added to label dead cells. Flow cytometry data were analyzed with FlowJo10.

### Statistical analysis

Statistical significance between mean values was determined using Student’s t-test or the Wilcoxon rank sum test. P-values for differences of proportions of clusters in scRNA-seq analysis were calculated using the Chi-square goodness of fit test. P-value correction in differential gene expression analysis with Seurat were performed with Bonferroni methods. P-values for BrdU decay kinetics experiment were calculated by simple linear regression in GraphPad Prism. ****: p < 0.0001; ***: p < 0.001; **: p < 0.01; *: p < 0.05. All outliers were included in the data.

## Ethics statement

All mice were maintained under pathogen-free conditions and handled in accordance with Guidelines of the Institutional Animal Care and Use Committee of RIKEN, Yokohama Branch (2018-075). Almost all available mice were used randomly for experiments without selection.

## Author Contributions

**Kenta Horie**, Data curation, Formal analysis, Funding acquisition, Investigation, Validation, Writing – original draft; **Kano Namiki**, Data curation, Formal analysis, Validation; **Maki Miyauchi,** Data curation; **Tatusya Ishikawa**, Investigation, Writing – review and editing, **Mio Hayama,** Validation; **Yuya Maruyama**; Validation, Writing – original draft; **Takahisa Miyao**, Supervision, Writing – review and editing; **Shigeo Murata,** Supervision, Writing – review and editing; **Nobuko Akiyama**, Formal analysis, Funding acquisition, Investigation, Supervision, Writing – review and editing; **Taishin Akiyama**, Formal analysis, Funding acquisition, Investigation, Project administration, Supervision, Validation, Writing – review and editing

## Acknowledgements

We thank the sequencing staff at the RIKEN Center for Integrative Medical Sciences for assisting with RNA-seq. Computations were performed on supercomputers at the National Institute of Genetics and at ISD, RIKEN (HOKUSAI).

---

**Supplementary Figure 1.**
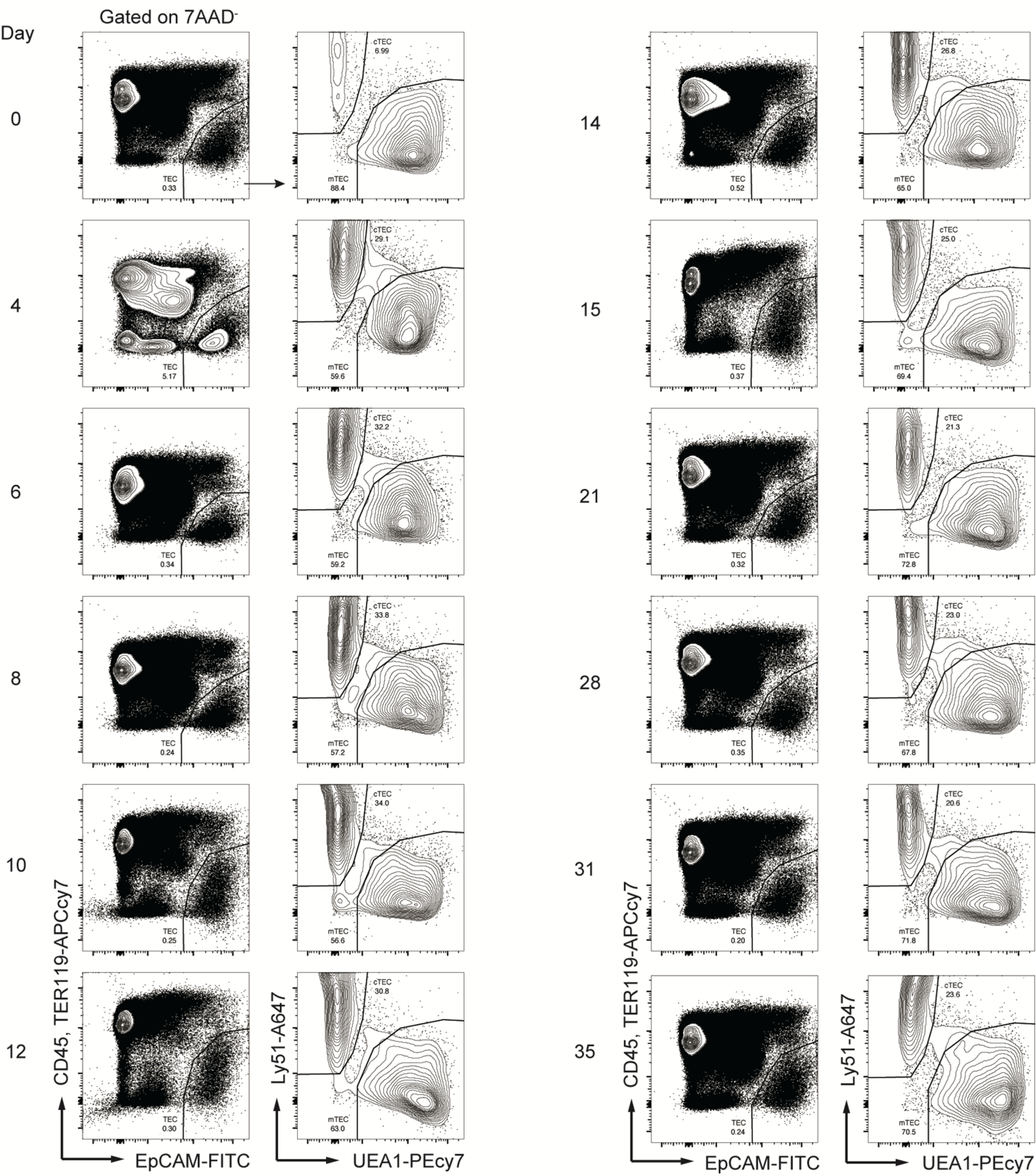
Flow cytometric analysis of the post-irradiation thymus Representative flow cytometer plots of TECs, each day after irradiation (related to figure1).

**Supplementary Figure 2.**
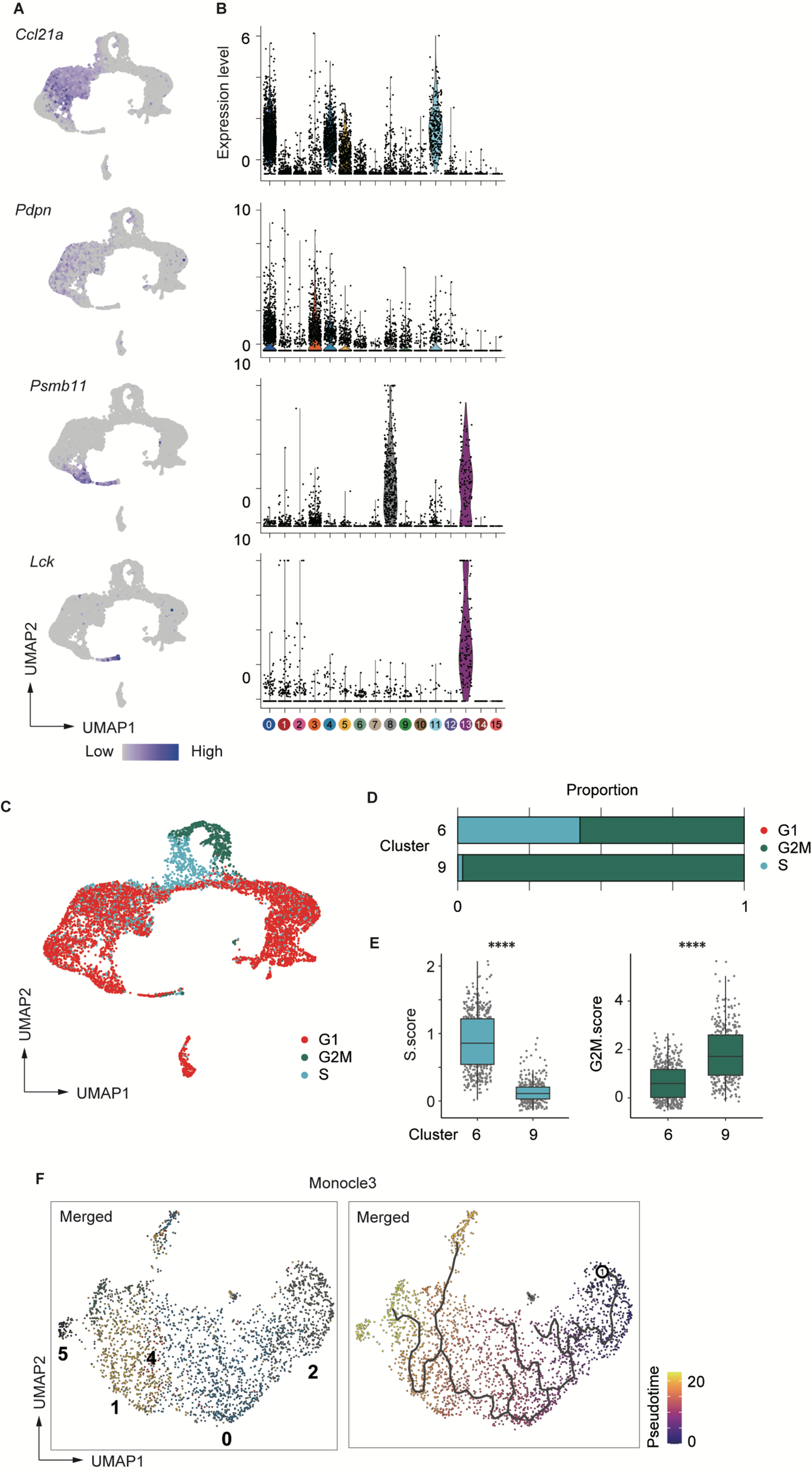
Analysis of scRNA-seq data. **(A)** Heatmaps of marker gene expression (counts per 10^4^ counts) projected onto UMAP plots. **(B)** Violin plots of marker gene expression (scaled counts per 10^4^ counts). **(C)** Cell-cycle scores of individual cells. All cells were assigned in G1, G2/M, and S phase of cell cycles based on expression of cell cycle related genes, using the Seurat package. **(D)** Cell-cycle scores of clusters 6 and 9 corresponding to TA-mTECs. **(E)** Scores of S (left) and G2/M (right) were plotted for individual cells from clusters 6 and 9. **(F)** Monocle 3 trajectory and pseudotime analysis of Aire+mTECs and Late-Aire mTEC subsets. ****: p < 0.0001; Wilcoxon rank sum test.

**Supplementary Figure 3.**
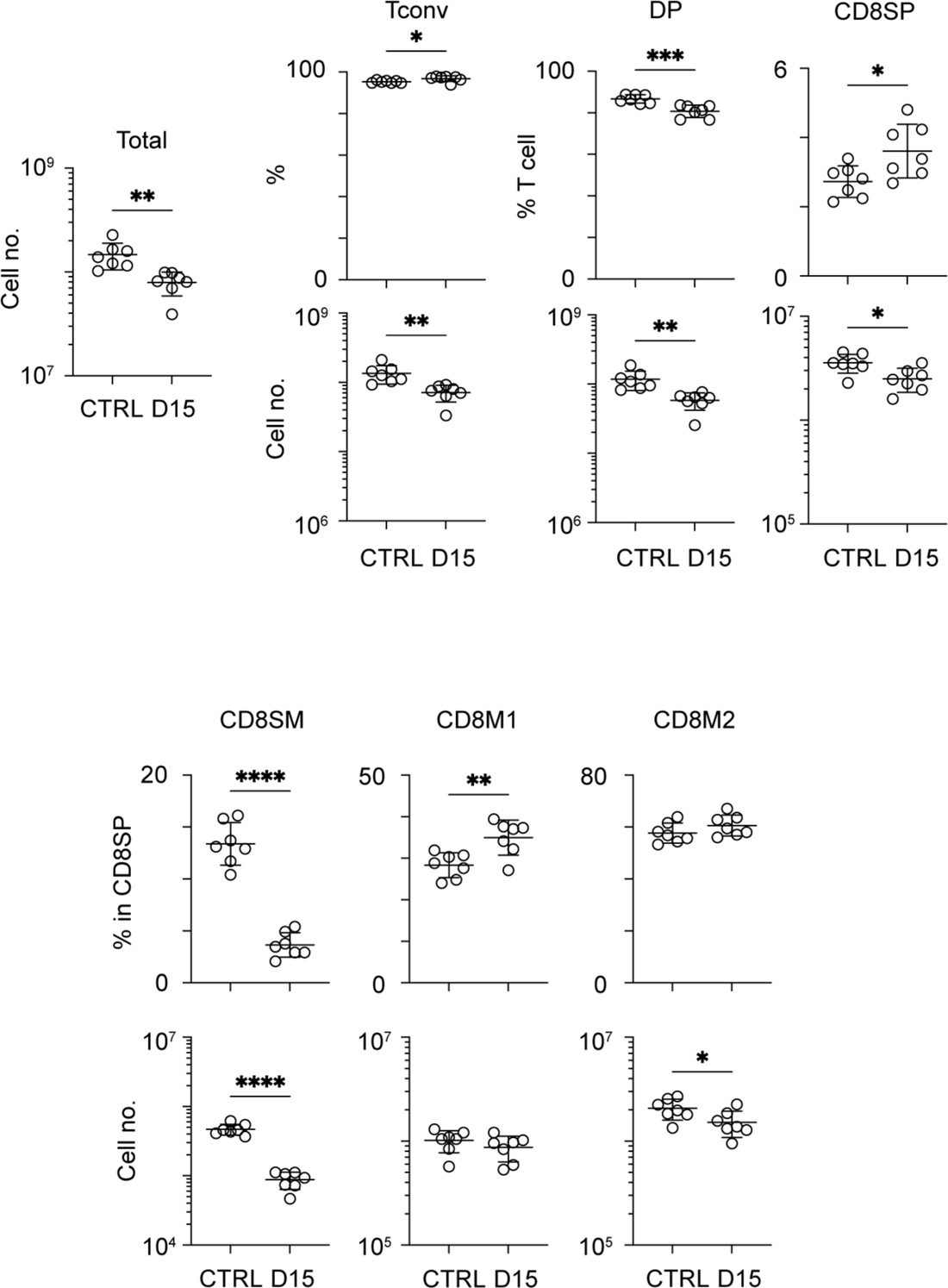
Flow-cytometric analysis of thymocytes. Summary data of flow-cytometric analysis (related to figure 6A., B). *: p < 0.05; **: p < 0.01; ***: p < 0.001; two-tailed t test.

## References

Abramson, J., and Anderson, G. (2017). Thymic Epithelial Cells. Annu Rev Immunol 35, 85–118. doi: 10.1146/annurev-immunol-051116-052320.

Akiyama, T., Shimo, Y., Yanai, H., Qin, J., Ohshima, D., Maruyama, Y., et al. (2008). The tumor necrosis factor family receptors RANK and CD40 cooperatively establish the thymic medullary microenvironment and self-tolerance. Immunity 29(3), 423–437. doi: 10.1016/j.immuni.2008.06.015.

Alawam, A.S., Cosway, E.J., James, K.D., Lucas, B., Bacon, A., Parnell, S.M., et al. (2022). Failures in thymus medulla regeneration during immune recovery cause tolerance loss and prime recipients for auto-GVHD. J Exp Med 219(2). doi: 10.1084/jem.20211239.

Anderson, M.S., Venanzi, E.S., Klein, L., Chen, Z., Berzins, S.P., Turley, S.J., et al. (2002). Projection of an immunological self shadow within the thymus by the aire protein. Science 298(5597), 1395–1401. doi: 10.1126/science.1075958.

Bornstein, C., Nevo, S., Giladi, A., Kadouri, N., Pouzolles, M., Gerbe, F., et al. (2018). Single-cell mapping of the thymic stroma identifies IL-25-producing tuft epithelial cells. Nature 559(7715), 622–626. doi: 10.1038/s41586-018-0346-1.

Bruserud, O., Oftedal, B.E., Wolff, A.B., and Husebye, E.S. (2016). AIRE-mutations and autoimmune disease. Curr Opin Immunol 43, 8–15. doi: 10.1016/j.coi.2016.07.003.

Cao, J., Spielmann, M., Qiu, X., Huang, X., Ibrahim, D.M., Hill, A.J., et al. (2019). The single-cell transcriptional landscape of mammalian organogenesis. Nature 566(7745), 496–502. doi: 10.1038/s41586-019-0969-x.

Derbinski, J., Schulte, A., Kyewski, B., and Klein, L. (2001). Promiscuous gene expression in medullary thymic epithelial cells mirrors the peripheral self. Nat Immunol 2(11), 1032–1039. doi: 10.1038/ni723.

Dhalla, F., Baran-Gale, J., Maio, S., Chappell, L., Hollander, G.A., and Ponting, C.P. (2020). Biologically indeterminate yet ordered promiscuous gene expression in single medullary thymic epithelial cells. EMBO J 39(1), e101828. doi: 10.15252/embj.2019101828.

Duah, M., Li, L., Shen, J., Lan, Q., Pan, B., and Xu, K. (2021). Thymus Degeneration and Regeneration. Front Immunol 12, 706244. doi: 10.3389/fimmu.2021.706244.

Dudakov, J.A., Hanash, A.M., Jenq, R.R., Young, L.F., Ghosh, A., Singer, N.V., et al. (2012). Interleukin-22 drives endogenous thymic regeneration in mice. Science 336(6077), 91–95. doi: 10.1126/science.1218004.

Ferreirinha, P., Ribeiro, C., Morimoto, J., Landry, J.J.M., Matsumoto, M., Meireles, C., et al. (2021). A novel method to identify Post-Aire stages of medullary thymic epithelial cell differentiation. Eur J Immunol 51(2), 311–318. doi: 10.1002/eji.202048764.

Gray, D., Abramson, J., Benoist, C., and Mathis, D. (2007). Proliferative arrest and rapid turnover of thymic epithelial cells expressing Aire. J Exp Med 204(11), 2521–2528. doi: 10.1084/jem.20070795.

Hauri-Hohl, M.M., Zuklys, S., Keller, M.P., Jeker, L.T., Barthlott, T., Moon, A.M., et al. (2008). TGF-beta signaling in thymic epithelial cells regulates thymic involution and postirradiation reconstitution. Blood 112(3), 626–634. doi: 10.1182/blood-2007-10-115618.

Hikosaka, Y., Nitta, T., Ohigashi, I., Yano, K., Ishimaru, N., Hayashi, Y., et al. (2008). The cytokine RANKL produced by positively selected thymocytes fosters medullary thymic epithelial cells that express autoimmune regulator. Immunity 29(3), 438–450. doi: 10.1016/j.immuni.2008.06.018.

Hsu, Y.C., Li, L., and Fuchs, E. (2014). Transit-amplifying cells orchestrate stem cell activity and tissue regeneration. Cell 157(4), 935–949. doi: 10.1016/j.cell.2014.02.057.

Irla, M., Hugues, S., Gill, J., Nitta, T., Hikosaka, Y., Williams, I.R., et al. (2008). Autoantigen-specific interactions with CD4+ thymocytes control mature medullary thymic epithelial cell cellularity. Immunity 29(3), 451–463. doi: 10.1016/j.immuni.2008.08.007.

Kaneko, K.B., Tateishi, R., Miyao, T., Takakura, Y., Akiyama, N., Yokota, R., et al. (2019). Quantitative analysis reveals reciprocal regulations underlying recovery dynamics of thymocytes and thymic environment in mice. Commun Biol 2, 444. doi: 10.1038/s42003-019-0688-8.

Kinsella, S., and Dudakov, J.A. (2020). When the Damage Is Done: Injury and Repair in Thymus Function. Front Immunol 11, 1745. doi: 10.3389/fimmu.2020.01745.

Lopes, N., Vachon, H., Marie, J., and Irla, M. (2017). Administration of RANKL boosts thymic regeneration upon bone marrow transplantation. EMBO Mol Med 9(6), 835–851. doi: 10.15252/emmm.201607176.

Michelson, D.A., Hase, K., Kaisho, T., Benoist, C., and Mathis, D. (2022). Thymic epithelial cells co-opt lineage-defining transcription factors to eliminate autoreactive T cells. Cell. doi: 10.1016/j.cell.2022.05.018.

Miller, C.N., Proekt, I., von Moltke, J., Wells, K.L., Rajpurkar, A.R., Wang, H., et al. (2018). Thymic tuft cells promote an IL-4-enriched medulla and shape thymocyte development. Nature 559(7715), 627–631. doi: 10.1038/s41586-018-0345-2.

Miyao, T., Miyauchi, M., Kelly, S.T., Terooatea, T.W., Ishikawa, T., Oh, E., et al. (2022). Integrative analysis of scRNA-seq and scATAC-seq revealed transit-amplifying thymic epithelial cells expressing autoimmune regulator. Elife 11. doi: 10.7554/eLife.73998.

Nishijima, H., Matsumoto, M., Morimoto, J., Hosomichi, K., Akiyama, N., Akiyama, T., et al. (2022). Aire Controls Heterogeneity of Medullary Thymic Epithelial Cells for the Expression of Self-Antigens. J Immunol 208(2), 303–320. doi: 10.4049/jimmunol.2100692.

Onder, L., Nindl, V., Scandella, E., Chai, Q., Cheng, H.W., Caviezel-Firner, S., et al. (2015). Alternative NF-kappa B signaling regulates mTEC differentiation from podoplanin-expressing presursors in the cortico-medullary junction. European Journal of Immunology 45(8), 2218–2231. doi: 10.1002/eji.201545677.

Park, J.E., Botting, R.A., Dominguez Conde, C., Popescu, D.M., Lavaert, M., Kunz, D.J., et al. (2020). A cell atlas of human thymic development defines T cell repertoire formation. Science 367(6480). doi: 10.1126/science.aay3224.

Rodrigues, P.M., Ribeiro, A.R., Perrod, C., Landry, J.J.M., Araujo, L., Pereira-Castro, I., et al. (2017). Thymic epithelial cells require p53 to support their long-term function in thymopoiesis in mice. Blood 130(4), 478–488. doi: 10.1182/blood-2016-12-758961.

Rossi, S.W., Jeker, L.T., Ueno, T., Kuse, S., Keller, M.P., Zuklys, S., et al. (2007a). Keratinocyte growth factor (KGF) enhances postnatal T-cell development via enhancements in proliferation and function of thymic epithelial cells. Blood 109(9), 3803–3811. doi: 10.1182/blood-2006-10-049767.

Rossi, S.W., Kim, M.Y., Leibbrandt, A., Parnell, S.M., Jenkinson, W.E., Glanville, S.H., et al. (2007b). RANK signals from CD4(+)3(-) inducer cells regulate development of Aire-expressing epithelial cells in the thymic medulla. J Exp Med 204(6), 1267–1272. doi: 10.1084/jem.20062497.

Sansom, S.N., Shikama-Dorn, N., Zhanybekova, S., Nusspaumer, G., Macaulay, I.C., Deadman, M.E., et al. (2014). Population and single-cell genomics reveal the Aire dependency, relief from Polycomb silencing, and distribution of self-antigen expression in thymic epithelia. Genome Res 24(12), 1918–1931. doi: 10.1101/gr.171645.113.

Schaue, D. (2017). A Century of Radiation Therapy and Adaptive Immunity. Front Immunol 8, 431. doi: 10.3389/fimmu.2017.00431.

Stuart, T., Butler, A., Hoffman, P., Hafemeister, C., Papalexi, E., Mauck, W.M., et al. (2019). Comprehensive Integration of Single-Cell Data. Cell 177(7), 1888-+. doi: 10.1016/j.cell.2019.05.031.

Wells, K.L., Miller, C.N., Gschwind, A.R., Wei, W., Phipps, J.D., Anderson, M.S., et al. (2020). Combined transient ablation and single cell RNA sequencing reveals the development of medullary thymic epithelial cells. Elife 9. doi: 10.7554/eLife.60188.

Wertheimer, T., Velardi, E., Tsai, J., Cooper, K., Xiao, S., Kloss, C.C., et al. (2018). Production of BMP4 by endothelial cells is crucial for endogenous thymic regeneration. Sci Immunol 3(19). doi: 10.1126/sciimmunol.aal2736.

Wurbel, M.A., Philippe, J.M., Nguyen, C., Victorero, G., Freeman, T., Wooding, P., et al. (2000). The chemokine TECK is expressed by thymic and intestinal epithelial cells and attracts double- and single-positive thymocytes expressing the TECK receptor CCR9. Eur J Immunol 30(1), 262–271. doi: 10.1002/1521-4141(200001)30:1<262::AID-IMMU262>3.0.CO;2-0.

Xing, Y., Wang, X., Jameson, S.C., and Hogquist, K.A. (2016). Late stages of T cell maturation in the thymus involve NF-kappaB and tonic type I interferon signaling. Nat Immunol 17(5), 565–573. doi: 10.1038/ni.3419.

Yahyapour, R., Amini, P., Rezapour, S., Cheki, M., Rezaeyan, A., Farhood, B., et al. (2018). Radiation-induced inflammation and autoimmune diseases. Mil Med Res 5(1), 9. doi: 10.1186/s40779-018-0156-7.

Zhou, W., Gao, F., Romero-Wolf, M., Jo, S., and Rothenberg, E.V. (2022). Single-cell deletion analyses show control of pro-T cell developmental speed and pathways by Tcf7, Spi1, Gata3, Bcl11a, Erg, and Bcl11b. Sci Immunol 7(71), eabm1920. doi: 10.1126/sciimmunol.abm1920.

